# Gut Microbiota Changes in Nonalcoholic Fatty Liver Disease and Concomitant Cardiovascular Diseases

**DOI:** 10.1101/2020.07.27.224329

**Authors:** Olena H. Kurinna

## Abstract

Nonalcoholic fatty liver disease (NAFLD) bears serious economic consequences for the health care system worldwide and Ukraine, in particular. Cardiovascular diseases (CVD) are the main cause of mortality in NAFLD patients. Changes in the gut microbiota composition can be regarded as a potential mechanism of CVD in NAFLD patients.

The purpose of this work was to investigate changes in major gut microbiota phylotypes, *Bacteroidetes, Firmicutes* and *Actinobacteria* with quantification of *Firmicutes/Bacteroidetes* in NAFLD patients with concomitant CVD.

The author enrolled 120 NAFLD subjects (25 with concomitant arterial hypertension (AH) and 24 with coronary artery disease (CAD)). The gut microbiota composition was assessed by qPCR.

**Results:** the author found a marked tendency towards an increase in the concentration of *Bacteroidetes* (by 37.11% and 21.30%, respectively) with a decrease in *Firmicutes* (by 7.38% and 7.77%, respectively) in both groups with comorbid CAD and AH with the identified changes not reaching a statistical significance. The author quantified a statistically significant decrease in the concentration of *Actinobacteria* in patients with NAFLD with concomitant CAD at 41.37% (p<0.05) as compared with those with an isolated NAFLD. In patients with concomitant AH, the content of *Actinobacteria* dropped by 12.35%, which was statistically insignificant.

**Conclusions:** the author established changes in the intestinal microbiota, namely decrease in *Actinobacteria* in patients with CAD, which requires further research.

## 1 Introduction

An excessive fat accumulation in hepatocytes with subsequent progressive inflammatory and fibrotic changes is a pathophysiological basis of most common liver pathological condition, i.e. non-alcoholic fatty liver disease (NAFLD)(1). In most cases, NAFLD shows a benign course, remaining at the stage of nonalcoholic steatosis (NAS), but in some cases, NAS progresses to a more severe stage, including nonalcoholic steatohepatitis (NASH), cirrhosis and even a hepatocellular carcinoma(2).

The significance of NAFLD in terms of public health is substantiated by its multifaceted impact on morbidity and mortality, as well as economic consequences for the health care system worldwide and Ukraine, in particular(3). According to the latest epidemiological data, 64 million NAFLD patients were identified in the United States alone, and 52 million in European countries(1, 4) which gives this pathology a ‘special weight’.

In addition to direct annual economic costs, the relevance of NAFLD is linked to the social consequences associated with a shorter life expectancy (adjusted for the life quality), and a burden of metabolic complications, including cardiovascular diseases (CVD). Furthermore, CVD are one of the leading causes of mortality among patients with NAFLD(1, 5).

An increase in a cardiovascular risk (CVR) in NAFLD patients is observable even in those with a relatively low body mass index (BMI)(6) and persists with an allowance for traditional risk factors, i.e. fatty liver and severe fibrosis as independent risk factors for CVD(7, 8).

Whereas the data on the role of intestinal microbiota in the development of atherosclerosis, hypertension and heart failure(9–11) are abandoned, the research in NAFLD patients with concomitant CVD is still scarce, and the relation between the microbial composition and the development of cardiovascular pathology in patients within this category remains unclear (12). Thus, further research in identifying holistic multifactorial microbiome-mediated mechanisms of NAFLD and CVD is required.(12)

## 3 Subjects and Methods

The study was carried out at the Department for the Study of GIS Diseases and their Comorbidity with Non-Communicable Diseases of the Governmental Institution “L.T. Malaya National Institute of Therapy of the National Academy of Medical Sciences of Ukraine” as part of the research project “To develop ways of individualized correction of metabolic disorders in patients with nonalcoholic fatty liver disease based on the study of intestinal microbiome, regulatory molecules and markers of systemic inflammation” (governmental registration № 017Γ003030). The study enrolled 120 NAFLD outpatients who represented the main group. The control group consisted of 9 healthy volunteers.

The study was approved of by the Local Ethics Committee of the Institution (Protocol No 04 of April, 2019 and 6 of July 17, 2020) as carried out in compliance with the Law of Ukraine “On Medical Products”, 1996, Art. 7, 8, 12; Principles of ICH GCP (2008); Ordinance of the Ministry of Health of Ukraine № 690 of September 23, 2009, “On Establishment of the Rules of Clinical Trials and Expert Examination of Clinical Trial Materials and the Standard Regulations on the Ethics Committee” with amendments. All patients signed an informed consent prior to the enrollment in the study and exposure to laboratory and instrumental procedures.

The patients examination was conducted in compliance with the recommendations of the European Association for the Study of the Liver (EASL) “EASL-EASD-EASO Clinical Practice Guidelines for the Management of Non-Alcoholic Fatty Liver Disease”(13); the national-adapted clinical guidelines “Non-alcoholic fatty liver disease” and “Non-alcoholic steatohepatitis”. All patients underwent an interview to determine secondary fatty liver etiologic factors, i.e. alcoholic abuse and drug-induced fatty liver. We also excluded viral hepatitis B and C, haemochromatosis, autoimmune hepatitis, coeliac disease, Wilson’s disease, etc. We did not enroll in this study patients with severe steatohepatitis and cirrhosis. Exclusion criteria included other conditions possibly affecting the composition of the gut microbiota, including bacterial overgrowth syndrome, antibiotic and probiotic treatment.

We evaluated the steatosis degree according to the NAS index, and fibrosis degree according to the METAVIR scale with the hepatobiliary system ultrasound visualization (Soneus P7 ultrasound scanning system) by the wave attenuation coefficient and shear wave elastometry, respectively. The value of the wave attenuation coefficient from 1 to 2.2 dB/cm indicated the absence of steatosis, the value from 2.2 to 2.3 dB/cm - the first degree steatosis, from 2.3 to 2.9 dB/cm – the second degree steatosis and from 2.9 to 3.5 dB/cm - the third degree steatosis. In turn, the value of the shear wave elastometry coefficient from 0 to 5.8 kPa indicated the absence of fibrosis, from 5.8 to 7.0 kPa – fibrosis FI, from 7.0 to 9.5 kPa – F II, from 9.5 to 12.5 kPa – F III, and more than 12.5 kPa was indicative of F4, or cirrhosis.

The CVD verification, in particular CAD and hypertension, was performed prior to the enrollment in the study, in compliance with ESC and ACC/AHA guidelines for the diagnosis and management of patients with stable coronary artery disease and arterial hypertension(14, 15). 25 (20.83%) patients in the main group were diagnosed as having concomitant arterial hypertension (AH), and 24 (20.00%) patients were diagnosed as those with CAD.

The assessment of the major gut microbiota phylotypes was performed by identifying the total bacterial DNA and DNA of *Bacteroidetes, Firmicutes* and *Actinobacteria* via the real-time quantitative polymerase chain reaction(16, 17). Firstly, we aliquoted freshly collected faeces samples in sterile containers with the following quick freezing and storage until extraction at - 20°C. Next, we extracted DNA from 400 mg of faeces with the use of a Ribo-prep nucleic acid extraction kit (AmpliSens, Russian Federation). We measured the DNA concentration in the extracts with the use of a Qubit 3 fluorometer with a Qubit dsDNA HS Assay Kits (Thermo Scientific, USA) and rectified it to ~ 10 ng/μl. Then we used the real-time PCR product detection system, CFX96 Touch (Bio-Rad, USA). Amplification program included:

– initial stage of denaturation for 5 min at 95°C – 40 cycles: 15 sec at 95°C, 15 sec at 61.5°C, 30 sec at 72°C with the reading of the fluorescence signal;
– the final stage of elongation - 5 min at 72°C.

We determined the anthropometric indices, i.e. height in metres; body weight in kilograms with the following quantification of the body mass index (BMI) and body composition with the use of bioimpedancemetry (OMRON BF 511, Japan, registration number № 20180102074). The distribution of adipose tissue was assessed by measuring the waist (WC) and hips (HC) circumferences and a quotient WC/HC. All patients were evaluated for their liver function, carbohydrate metabolism and lipid metabolism. We assessed liver function by measuring the alanine and asparagine transaminases, alkaline phosphatase and gamma-glutamyl transferase (automatic biochemical analyzer “HumaStar 200”, registration number № 21150707005, Germany). We evaluated carbohydrate metabolism through the serum glucose concentration (automatic biochemical analyzer “HumaStar 200”, registration number № 21150707005, Germany), glycosylated hemoglobin (automatic biochemical analyzer photometer “Humalyzer 2000”, registration number № 18300, Germany), insulin (DRG Instruments GmbH, Germany) followed by a quantitative assessment of insulin resistance according to the HOMA-IR. As proinflammatory markers we assessed serum C-reactive protein («CRP-BEST», LLC Best Diagnostic, Ukraine) and tumor necrosis α concentrations («EIA-TNF-alpha», Cytokine, Russian Federation).

To exclude the bacterial overgrowth syndrome, we used the hydrogen breath test (gas analyzer Gastro + Gastrolyzer, № 12-8989, UK).

The statistical analysis was performed with the use the software package ‘STATISTICA 13.1’ (Statsoft, USA). According to the Kolmogorov-Smirnov criterion, the distribution of all studied parameters was established as different from normal (Gaussian), so we used non-parametric statistical methods for the data processment, hereinafter referred to as Me [LQ; UQ], where Me is the median, and LQ and UQ are the lower and upper quartiles, respectively. We used the Kraskel-Wallace test and the median test to investigate the dependence of the variables on the groups.

## 4 Results

### 4.1 Clinical characteristics

Clinical characteristics of patients are shown in Table 1.

**Table 1.**
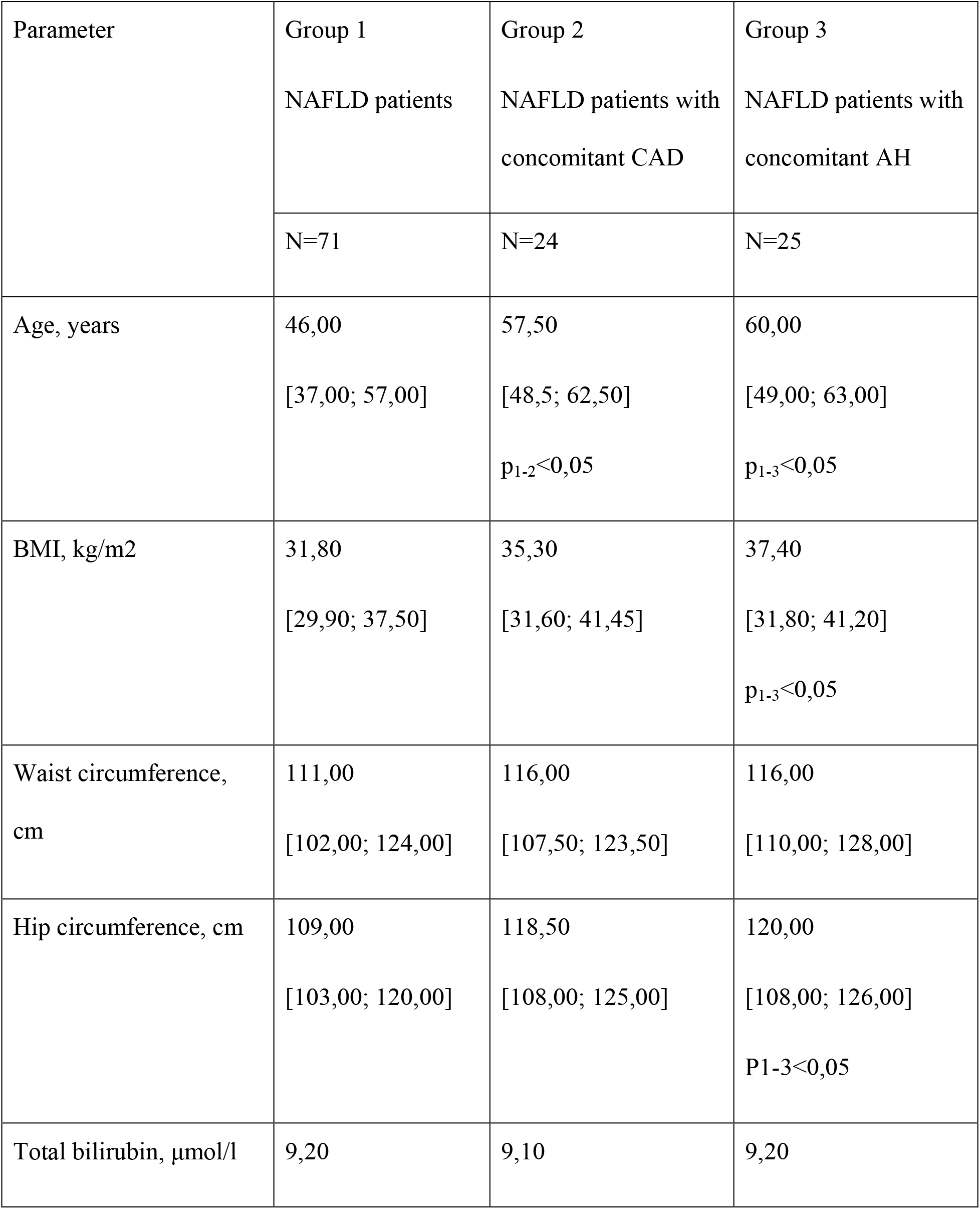
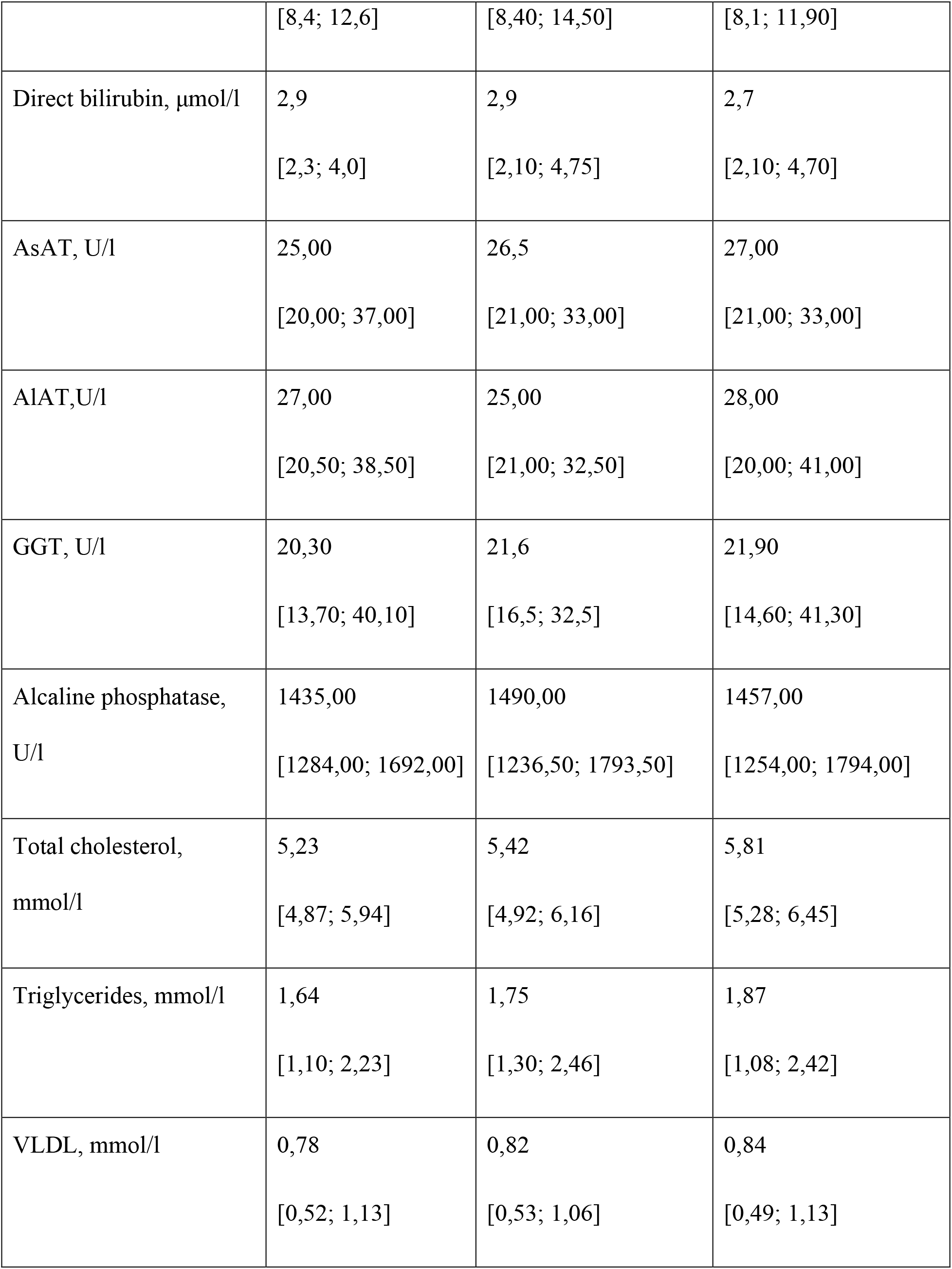
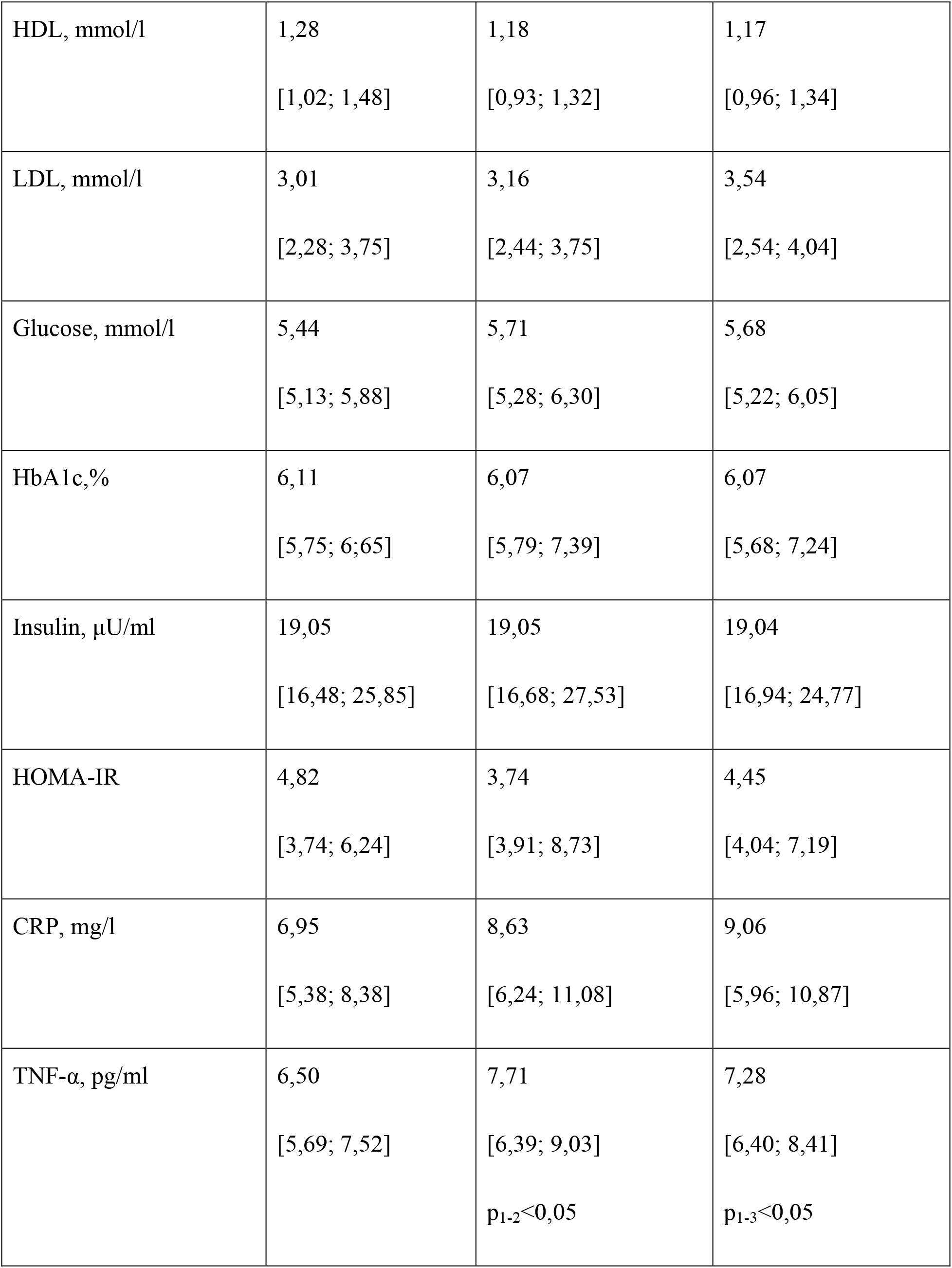
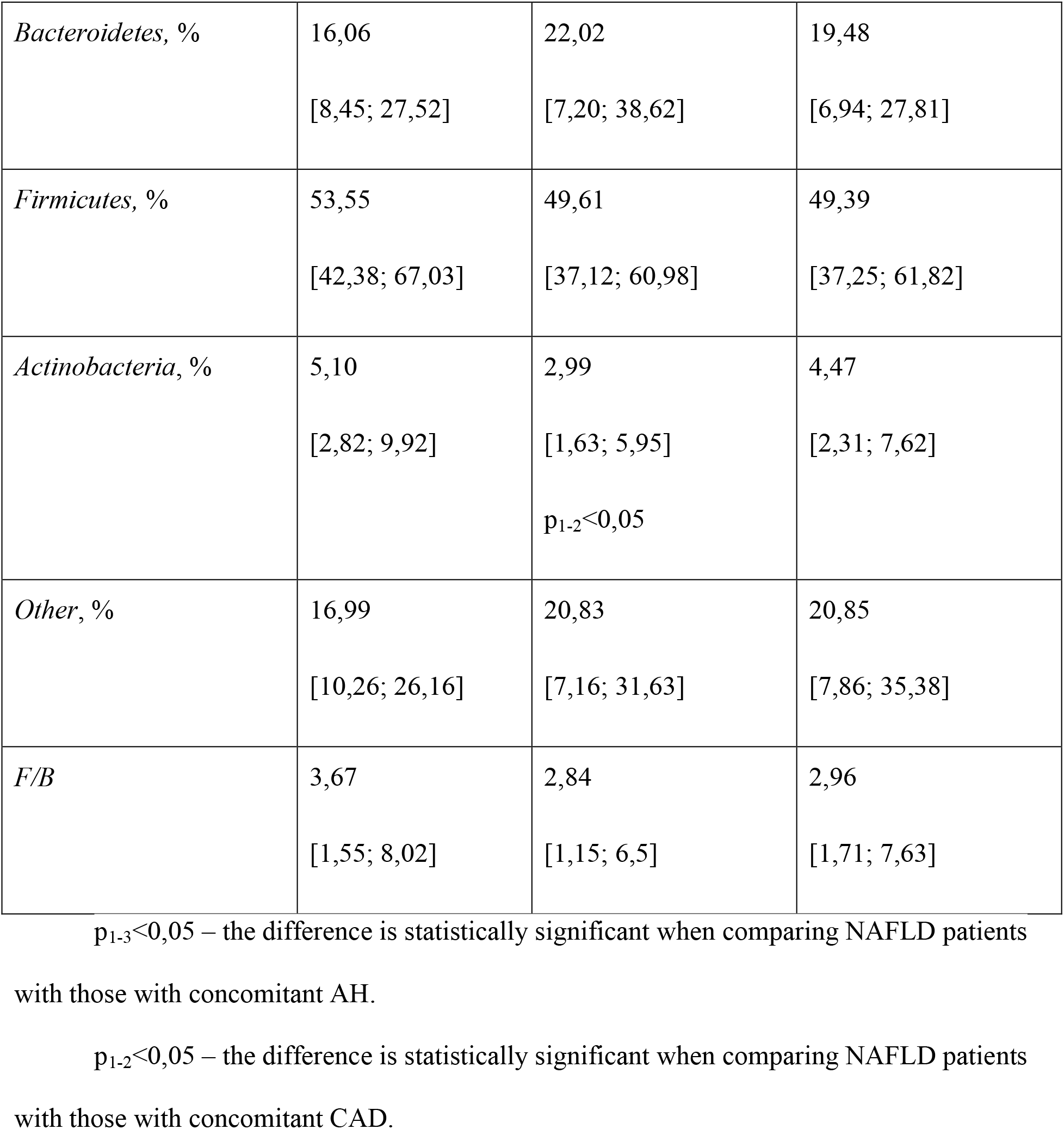
Clinical characteristics of the examined patients

| Parameter | Group 1 | Group 2 | Group 3 |
| --- | --- | --- | --- |
|  | NAFLD patients | NAFLD patients with concomitant CAD | NAFLD patients with concomitant AH |
|  | N=71 | N=24 | N=25 |
| Age, years | 46,00<br>[37,00; 57,00] | 57,50<br>[48,5; 62,50]<br>$p_{1-2}<0,05$ | 60,00<br>[49,00; 63,00]<br>$p_{1-3}<0,05$ |
| BMI, kg/m<sup>2</sup> | 31,80<br>[29,90; 37,50] | 35,30<br>[31,60; 41,45] | 37,40<br>[31,80; 41,20]<br>$p_{1-3}<0,05$ |
| Waist circumference, cm | 111,00<br>[102,00; 124,00] | 116,00<br>[107,50; 123,50] | 116,00<br>[110,00; 128,00] |
| Hip circumference, cm | 109,00<br>[103,00; 120,00] | 118,50<br>[108,00; 125,00] | 120,00<br>[108,00; 126,00]<br>$P_{1-3}<0,05$ |
| Total bilirubin, $\mu\text{mol/l}$ | 9,20 | 9,10 | 9,20 |
|  | [8,4; 12,6] | [8,40; 14,50] | [8,1; 11,90] |
| Direct bilirubin, $\mu\text{mol/l}$ | 2,9<br>[2,3; 4,0] | 2,9<br>[2,10; 4,75] | 2,7<br>[2,10; 4,70] |
| AsAT, U/l | 25,00<br>[20,00; 37,00] | 26,5<br>[21,00; 33,00] | 27,00<br>[21,00; 33,00] |
| AlAT, U/l | 27,00<br>[20,50; 38,50] | 25,00<br>[21,00; 32,50] | 28,00<br>[20,00; 41,00] |
| GGT, U/l | 20,30<br>[13,70; 40,10] | 21,6<br>[16,5; 32,5] | 21,90<br>[14,60; 41,30] |
| Alcaline phosphatase,<br>U/l | 1435,00<br>[1284,00; 1692,00] | 1490,00<br>[1236,50; 1793,50] | 1457,00<br>[1254,00; 1794,00] |
| Total cholesterol,<br>mmol/l | 5,23<br>[4,87; 5,94] | 5,42<br>[4,92; 6,16] | 5,81<br>[5,28; 6,45] |
| Triglycerides, mmol/l | 1,64<br>[1,10; 2,23] | 1,75<br>[1,30; 2,46] | 1,87<br>[1,08; 2,42] |
| VLDL, mmol/l | 0,78<br>[0,52; 1,13] | 0,82<br>[0,53; 1,06] | 0,84<br>[0,49; 1,13] |
| HDL, mmol/l | 1,28<br>[1,02; 1,48] | 1,18<br>[0,93; 1,32] | 1,17<br>[0,96; 1,34] |
| LDL, mmol/l | 3,01<br>[2,28; 3,75] | 3,16<br>[2,44; 3,75] | 3,54<br>[2,54; 4,04] |
| Glucose, mmol/l | 5,44<br>[5,13; 5,88] | 5,71<br>[5,28; 6,30] | 5,68<br>[5,22; 6,05] |
| HbA1c, % | 6,11<br>[5,75; 6,65] | 6,07<br>[5,79; 7,39] | 6,07<br>[5,68; 7,24] |
| Insulin, μU/ml | 19,05<br>[16,48; 25,85] | 19,05<br>[16,68; 27,53] | 19,04<br>[16,94; 24,77] |
| HOMA-IR | 4,82<br>[3,74; 6,24] | 3,74<br>[3,91; 8,73] | 4,45<br>[4,04; 7,19] |
| CRP, mg/l | 6,95<br>[5,38; 8,38] | 8,63<br>[6,24; 11,08] | 9,06<br>[5,96; 10,87] |
| TNF-α, pg/ml | 6,50<br>[5,69; 7,52] | 7,71<br>[6,39; 9,03]<br>$p_{1-2} < 0,05$ | 7,28<br>[6,40; 8,41]<br>$p_{1-3} < 0,05$ |
| *Bacteroidetes*, % | 16,06<br>[8,45; 27,52] | 22,02<br>[7,20; 38,62] | 19,48<br>[6,94; 27,81] |
| *Firmicutes*, % | 53,55<br>[42,38; 67,03] | 49,61<br>[37,12; 60,98] | 49,39<br>[37,25; 61,82] |
| *Actinobacteria*, % | 5,10<br>[2,82; 9,92] | 2,99<br>[1,63; 5,95]<br>$p_{1-2}<0,05$ | 4,47<br>[2,31; 7,62] |
| *Other*, % | 16,99<br>[10,26; 26,16] | 20,83<br>[7,16; 31,63] | 20,85<br>[7,86; 35,38] |
| *F/B* | 3,67<br>[1,55; 8,02] | 2,84<br>[1,15; 6,5] | 2,96<br>[1,71; 7,63] |

$p_{1-3}<0,05$ – the difference is statistically significant when comparing NAFLD patients with those with concomitant AH.
$p_{1-2}<0,05$ – the difference is statistically significant when comparing NAFLD patients with those with concomitant CAD.

The patient clinical examination showed an increase in BMI in all groups, with a statistical significance determined only in the AH group, in which BMI exceeded this parameter in CAD and isolated NAFLD patients by 4.82% and 17.61% (p<0.05), respectively. Similarly, the AH group displayed a statistically significant increase in HC, i.e. by 10.09%, compared with isolated NAFLD patients (p<0.05). The CAD patients revealed a similar HC tendency, which, however, did not reach the statistical significance (p=0.08).

All patients displayed no statistically significant changes in liver function, lipid and carbohydrate metabolism, which could be explained through the adequate therapy administered upon the case diagnosis. In particular, the cholesterol target values during the statin treatment were achieved in 8 patients (30.00%).

We observed an increase in the concentration of pro-inflammatory factors in both, AH and CAD, groups. Compared with isolated NAFLD patients, the CRP concentration proved an increase in CAD patients by 24.17%, and by 30.36% in AH patients, without reaching any statistical significance (p=0.05 and p=0.06, respectively). However, similar changes in the TNF-α concentration with an elevation by 18.62% in CAD patients and by 12.00% in the AH group proved statistically significant (p<0.05 for both groups).

### 4.2 Gut microbiota changes

The present study in the gut microbiota composition showed that, in comparison with isolated NAFLD cases, both CAD and AH groups developed a marked tendency towards an increase in the concentration of *Bacteroidetes* (by 37.11% and 21.30%, respectively) with a decrease in *Firmicutes* (by 7.38% and 7.77%, respectively) without reaching, however, any statistical significance. Conversely, the *F/B* ratio reached the maximum value in patients with isolated NAFLD with a decrease in CAD and AH groups by 22.62% and 19.35%, respectively.

On the other hand, we discovered a statistically significant drop in the *Actinobacteria* concentration only in CAD group (by 41.37%, p<0.05) whereas AH patients displayed an insignificant decrease by 12.35%.

Thus, we established anthropometric disorders, proinflammatory abnormalities in NAFLD patients with concomitant CVD and changes in the gut microbiota composition, all of which require particular research.

We investigated the interdependence between these changes and the medication treatment and found that the patients who reached the target values of LDL in the course of statin treatment showed an increase in *Bacteroidetes* by 1.90 times (p<0.05) with a decrease in the *F/B* ratio by 2.7 times (p<0.05).

These findings considered, we developed a mnemonic positional code to study the multifactorial relation between the treatment prescribed and gut microbiota changes, which enabled to prove deviation only for *Actinobacteria* concentration (H = 7.14, p = 0.06).

## 5 Discussion

The aim of this study was to investigate changes in the gut microbiota composition in NAFLD and CVD (comorbid in most cases).

As is known, intestinal microorganisms form a community, the gut microbiota, a diverse ecosystem within the human body, which combines about 10^14^ microorganisms(7). The vast majority of gut bacteria fall in the main five phylotypes, namely *Bacteroidetes* (56%), *Firmicutes* (29%), *Actinobacteria* (6%) and *Proteobacteria* (4%)(7). The gut microbiota can be called an ‘invisible organ’,(18) whose harmonious functioning is important for maintaining the health of the human body, whereas disturbances in the qualitative and quantitative composition of gut microbiota contribute to the development of various conditions, including obesity(19), type 2 diabetes mellitus(20), NAFLD (21) and CVD(22). The latter are known to be the leading cause of morbidity among NAFLD patients worldwide(23). The nature of the pathogenetic links between the gut microbiota changes in CVD in NAFLD patients still remains the subject of scientific research.

As is known, atherosclerosis is the key pathogenetic mechanism of CVD evolvement and progression.(24) Among the factors of atherogenesis, gut microorganisms are believed to play an important if not the leading role.(25–27) Cui L. et al. discovered changes in the quantitative composition of the gut microbiota at the phylotype level, e.g. a decrease in *Bacteroidetes* with a reciprocal increase in *Firmicutes*.(28) Despite the unfavorable results of the meta-analysis of the antibiotic effectiveness in CAD patients,(29) recent evidence demonstrated that the gut microbiota plays a causal role in atherosclerosis due to intestinal barrier dysfunction, endotoxemia, proinflammatory changes, the synthesis of trimethylamine N-oxide, bile acids and microbial metabolites(30), as well as adipose tissue dysfunction, disorders in lipid(31) and carbohydrate metabolism(32). In addition, microorganisms can realize their influence through the synthesis of short-chain fatty acids, butyrate in particular – the fermentation end products of dietary fiber, which are the main source of energy for colonocytes that support the intestinal mucosal barrier(33).

However, our results did not identify significant changes in *Bacteroidetes* and *Firmicutes*, despite certain visible tendencies. On the contrary, we found a statistically significant decrease in the *Actinobacteria* concentration. The study of open sources, however, shows that *Actinobacteria* mostly remain beyond scientific research.

*Actinobacteria* are microorganisms that share characteristics of both, bacteria and fungi, and are widespread in both terrestrial and aquatic ecosystems, while accounting only for 6% in human gut microbiota(34). The class *Actinobacteria* is further divided into 16 orders, the most investigated of which, due to its role in atherogenesis, is *Bifidobacteriales*.

Contrary to the preliminary data on the increased amount of *Actinobacteria* in the structure of atherosclerotic plaque(35), the results of the Tampere Sudden Death Study did not confirm this but showed that despite the tendency towards reduction in *Bifidobacterium* spp., the main representative of *Actinobacteria*, in CAD patients there was an inverse relation between the quantity of these bacteria and the severity of fibrosis, which suggests a favorable role of *Bifidobacterium spp*. in the prevention of coronary atherosclerosis(36). Potential mechanisms that account for these effects include the ability of *Bifidobacteria* to improve the structural and functional state of the intestine, which leads to inhibition of bacterial translocation and endotoxinemia reduction(36).

Changes in lipid metabolism is another phenomenon associated with gut microbiota. In particular, after 12 weeks of high fat diet in ApoE -/- mice, Li L. et al. observed a significant increase in the of quantity of *Firmicutes* with a decrease in *Actinobacteria, Bacteroidetes and Verrucomicrobia*(37). In addition, the authors discovered a negative correlation between the phylotype of *Actinobacteria* and the index of hyperlipidemia, weight gain, relative amount of epididymal fat and liver weight(37).

Other clinical studies showed that the quantity of *Actinobacteria* depends on physical activity(38). In particular, endurance exercises and/or cardiac exercise significantly increase the concentration of these microorganisms in the intestinal content.(38) Conversely, a sedentary lifestyle is a risk factor for both NAFLD and CAD.(39, 40)

## Conclusions

Our study supported the view that NAFLD is a socially and economically significant disease, most often with a comorbid course, especially with CVD.

This work showed that NAFLD patients with concomitant CVD, in particular, CAD and AH, developed significant violations of anthropometric parameters and pro-inflammatory markers. We observed no noticeable changes in liver function, lipid and carbohydrate metabolism with all group of patients, which can be explained through the appropriate therapeutic measures.

The research in the gut microbiota composition showed statistically significant changes in patients with NAFLD and concomitant CAD, i.e. a decrease in the concentration of *Actinobacteria* by 41.37% (p<0.05), compared with the patients with isolated NAFLD. However, the dynamics of the other phylotypes, *Bacteroidetes* and *Firmicutes*, were only tendencies.

These changes can be caused by malnutrition and reduced physical activity. All these suggest the potential protective role of the *Actinobacteria* phylotype in inhibiting the development of CAD, which justifies further research.

## Structured disclosures

### Funding/Support

This research was funded by the Government as part of the research project “To develop ways of individualized correction of metabolic disorders in patients with nonalcoholic fatty liver disease based on the study of intestinal microbiome, regulatory molecules and markers of systemic inflammation” (Governmental Registration № 017Γ003030).

### Disclosures

The author has declared no financial relationships with any organizations that might have an interest in the submitted work; nor any other relationships or activities that could appear to have influenced the submitted work.

### Ethical approval

All procedures in this study were performed in accordance with the ethical standards of the institutional research committee and the local ethical committee, and with the 1964 Helsinki Declaration and its later amendments. This research was approved of by the Local Ethical Committee: protocol reference number 4 of 10 April 2019. Before the enrollment in this research all patients signed the informed consent.

### Disclaimer

The author of this article is solely responsible for the content thereof; the publication of the article shall not constitute nor be deemed to constitute any representation by the Editors that the data presented therein are correct or sufficient to support the conclusions reached or that the experiment design or methodology is adequate.

### Previous presentations

None.

## Acknowledgments

The author wishes to thank Prof. Galyna D. Fadieienko, Director of the Governmental Institution “L.T. Malaya National Institute of the National Academy of Medical Sciences of Ukraine” for her expert advice and professional consultations. The author is also thankful to Valentyna Yu. Galchinskaya, Head of the Laboratory of Biochemical and Immune Enzyme Methods with Morphology, for her assistance in laboratory testing. The author feels indebted to Igor V. Ilyin for professional editing service.

## The author’s contributions

conceived and designed the analysis; collected the data, created the data pool and performed the analysis; wrote the paper.

## References

1. Mitra S, De A, Chowdhury A. 2020. Epidemiology of non-alcoholic and alcoholic fatty liver diseases. Transl Gastroenterol Hepatol 5:16.

2. Nadasdi A, Somogyi A, Igaz P, Firneisz G. 2018. [Non-alcoholic fatty liver disease - a summary and update based on the EASL-EASD-EASO Clinical Practice Guidelines of 2016]. Orv Hetil 159:1815–1830.

3. Rinella ME. 2015. Nonalcoholic fatty liver disease: a systematic review. Jama 313:2263–73.

4. Asrani SK, Devarbhavi H, Eaton J, Kamath PS. 2019. Burden of liver diseases in the world. J Hepatol 70:151–171.

5. Stepanova M, Rafiq N, Makhlouf H, Agrawal R, Kaur I, Younoszai Z, McCullough A, Goodman Z, Younossi ZM. 2013. Predictors of all-cause mortality and liver-related mortality in patients with non-alcoholic fatty liver disease (NAFLD). Dig Dis Sci 58:3017–23.

6. Barik A, Shah RV, Spahillari A, Murthy VL, Ambale-Venkatesh B, Rai RK, Das K, Santra A, Hembram JR, Bhattacharya D, Freedman JE, Lima J, Das R, Bhattacharyya P, Das S, Chowdhury A. 2016. Hepatic steatosis is associated with cardiometabolic risk in a rural Indian population: A prospective cohort study. Int J Cardiol 225:161–166.

7. Ma J, Li H. 2018. The Role of Gut Microbiota in Atherosclerosis and Hypertension. Front Pharmacol 9:1082.

8. Tana C, Ballestri S, Ricci F, Di Vincenzo A, Ticinesi A, Gallina S, Giamberardino MA, Cipollone F, Sutton R, Vettor R, Fedorowski A, Meschi T. 2019. Cardiovascular Risk in Non-Alcoholic Fatty Liver Disease: Mechanisms and Therapeutic Implications. Int J Environ Res Public Health 16.

9. Karbach SH, Schonfelder T, Brandao I, Wilms E, Hormann N, Jackel S, Schuler R, Finger S, Knorr M, Lagrange J, Brandt M, Waisman A, Kossmann S, Schafer K, Munzel T, Reinhardt C, Wenzel P. 2016. Gut Microbiota Promote Angiotensin II-Induced Arterial Hypertension and Vascular Dysfunction. J Am Heart Assoc 5.

10. Kim S, Goel R, Kumar A, Qi Y, Lobaton G, Hosaka K, Mohammed M, Handberg EM, Richards EM, Pepine CJ, Raizada MK. 2018. Imbalance of gut microbiome and intestinal epithelial barrier dysfunction in patients with high blood pressure. Clin Sci (Lond) 132:701–718.

11. Tang WHW, Li DY, Hazen SL. 2019. Dietary metabolism, the gut microbiome, and heart failure. Nat Rev Cardiol 16:137–154.

12. Kazemian N, Mahmoudi M, Halperin F, Wu JC, Pakpour S. 2020. Gut microbiota and cardiovascular disease: opportunities and challenges. Microbiome 8:36.

13. Marchesini G. 2016. EASL-EASD-EASO Clinical Practice Guidelines for the management of non-alcoholic fatty liver disease. J Hepatol 64:1388–402.

14. Joseph J, Velasco A, Hage FG, Reyes E. 2018. Guidelines in review: Comparison of ESC and ACC/AHA guidelines for the diagnosis and management of patients with stable coronary artery disease. J Nucl Cardiol 25:509–515.

15. Williams B, Mancia G, Spiering W, Agabiti Rosei E, Azizi M, Burnier M, Clement DL, Coca A, de Simone G, Dominiczak A, Kahan T, Mahfoud F, Redon J, Ruilope L, Zanchetti A, Kerins M, Kjeldsen SE, Kreutz R, Laurent S, Lip GYH, McManus R, Narkiewicz K, Ruschitzka F, Schmieder RE, Shlyakhto E, Tsioufis C, Aboyans V, Desormais I. 2018. 2018 ESC/ESH Guidelines for the management of arterial hypertension. Eur Heart J 39:3021–3104.

16. Bacchetti De Gregoris T, Aldred N, Clare AS, Burgess JG. 2011. Improvement of phylum- and class-specific primers for real-time PCR quantification of bacterial taxa. J Microbiol Methods 86:351–6.

17. Turnbaugh PJ, Hamady M, Yatsunenko T, Cantarel BL, Duncan A, Ley RE, Sogin ML, Jones WJ, Roe BA, Affourtit JP, Egholm M, Henrissat B, Heath AC, Knight R, Gordon JI. 2009. A core gut microbiome in obese and lean twins. Nature 457:480–4.

18. Li X, Liu L, Cao Z, Li W, Li H, Lu C, Yang X, Liu Y. 2020. Gut microbiota as an “invisible organ” that modulates the function of drugs. Biomedicine & pharmacotherapy = Biomedecine & pharmacotherapie 121:109653.

19. John GK, Mullin GE. 2016. The Gut Microbiome and Obesity. Curr Oncol Rep 18:45.

20. Pedersen HK, Gudmundsdottir V, Nielsen HB, Hyotylainen T, Nielsen T, Jensen BA, Forslund K, Hildebrand F, Prifti E, Falony G, Le Chatelier E, Levenez F, Dore J, Mattila I, Plichta DR, Poho P, Hellgren LI, Arumugam M, Sunagawa S, Vieira-Silva S, Jorgensen T, Holm JB, Trost K, Kristiansen K, Brix S, Raes J, Wang J, Hansen T, Bork P, Brunak S, Oresic M, Ehrlich SD, Pedersen O. 2016. Human gut microbes impact host serum metabolome and insulin sensitivity. Nature 535:376–81.

21. Kolodziejczyk AA, Zheng D, Shibolet O, Elinav E. 2019. The role of the microbiome in NAFLD and NASH. EMBO Mol Med 11.

22. Emoto T, Yamashita T, Kobayashi T, Sasaki N, Hirota Y, Hayashi T, So A, Kasahara K, Yodoi K, Matsumoto T, Mizoguchi T, Ogawa W, Hirata KI. 2017. Characterization of gut microbiota profiles in coronary artery disease patients using data mining analysis of terminal restriction fragment length polymorphism: gut microbiota could be a diagnostic marker of coronary artery disease. Heart Vessels 32:39–46.

23. Liu Y, Zhong GC, Tan HY, Hao FB, Hu JJ. 2019. Nonalcoholic fatty liver disease and mortality from all causes, cardiovascular disease, and cancer: a meta-analysis. Sci Rep 9:11124.

24. Gui T, Shimokado A, Sun Y, Akasaka T, Muragaki Y. 2012. Diverse roles of macrophages in atherosclerosis: from inflammatory biology to biomarker discovery. Mediators Inflamm 2012:693083.

25. Drosos I, Tavridou A, Kolios G. 2015. New aspects on the metabolic role of intestinal microbiota in the development of atherosclerosis. Metabolism 64:476–81.

26. Gregory JC, Buffa JA, Org E, Wang Z, Levison BS, Zhu W, Wagner MA, Bennett BJ, Li L, DiDonato JA, Lusis AJ, Hazen SL. 2015. Transmission of atherosclerosis susceptibility with gut microbial transplantation. J Biol Chem 290:5647–60.

27. Jie Z, Xia H, Zhong SL, Feng Q, Li S, Liang S, Zhong H, Liu Z, Gao Y, Zhao H, Zhang D, Su Z, Fang Z, Lan Z, Li J, Xiao L, Li J, Li R, Li X, Li F, Ren H, Huang Y, Peng Y, Li G, Wen B, Dong B, Chen JY, Geng QS, Zhang ZW, Yang H, Wang J, Wang J, Zhang X, Madsen L, Brix S, Ning G, Xu X, Liu X, Hou Y, Jia H, He K, Kristiansen K. 2017. The gut microbiome in atherosclerotic cardiovascular disease. Nat Commun 8:845.

28. Cui L, Zhao T, Hu H, Zhang W, Hua X. 2017. Association Study of Gut Flora in Coronary Heart Disease through High-Throughput Sequencing. Biomed Res Int 2017:3796359.

29. Andraws R, Berger JS, Brown DL. 2005. Effects of antibiotic therapy on outcomes of patients with coronary artery disease: a meta-analysis of randomized controlled trials. Jama 293:2641–7.

30. Kasahara K, Tanoue T, Yamashita T, Yodoi K, Matsumoto T, Emoto T, Mizoguchi T, Hayashi T, Kitano N, Sasaki N, Atarashi K, Honda K, Hirata KI. 2017. Commensal bacteria at the crossroad between cholesterol homeostasis and chronic inflammation in atherosclerosis. J Lipid Res 58:519–528.

31. Schoeler M, Caesar R. 2019. Dietary lipids, gut microbiota and lipid metabolism. Rev Endocr Metab Disord 20:461–472.

32. Ganesan K, Chung SK, Vanamala J, Xu B. 2018. Causal Relationship between Diet-Induced Gut Microbiota Changes and Diabetes: A Novel Strategy to Transplant Faecalibacterium prausnitzii in Preventing Diabetes. Int J Mol Sci 19.

33. Trøseid M, Andersen G, Broch K, Hov JR. 2020. The gut microbiome in coronary artery disease and heart failure: Current knowledge and future directions. EBioMedicine 52:102649.

34. Anandan R, Dharumadurai D, Manogaran GP. 2016. Actinobacteria - Basics and Biotechnological Applications. In Dhanasekaran D, Jiang Y (ed), An Introduction to Actinobacteria. IntechOpen, doi:DOI: 10.5772/62329.. https://www.intechopen.com/books/actinobacteria-basics-and-biotechnological-applications/an-introduction-to-actinobacteria.

35. Lindskog Jonsson A, Hallenius FF, Akrami R, Johansson E, Wester P, Arnerlov C, Backhed F, Bergstrom G. 2017. Bacterial profile in human atherosclerotic plaques. Atherosclerosis 263:177–183.

36. Tuomisto S, Huhtala H, Martiskainen M, Goebeler S, Lehtimäki T, Karhunen PJ. 2019. Age-dependent association of gut bacteria with coronary atherosclerosis: Tampere Sudden Death Study. PLoS One 14:e0221345.

37. Li L, Shi M, Salerno S, Tang M, Guo F, Liu J, Feng Y, Fu M, Huang Q, Ma L, Li Y, Fu P. 2019. Microbial and metabolomic remodeling by a formula of Sichuan dark tea improves hyperlipidemia in apoE-deficient mice. PLoS One 14:e0219010.

38. Mach N, Fuster-Botella D. 2017. Endurance exercise and gut microbiota: A review. J Sport Health Sci 6:179–197.

39. Kim D, Vazquez-Montesino LM, Li AA, Cholankeril G, Ahmed A. Inadequate Physical Activity and Sedentary Behavior Are Independent Predictors of Nonalcoholic Fatty Liver Disease. Hepatology n/a.

40. Zhuang Z, Gao M, Yang R, Li N, Liu Z, Cao W, Huang T. 2020. Association of physical activity, sedentary behaviours and sleep duration with cardiovascular diseases and lipid profiles: a Mendelian randomization analysis. Lipids Health Dis 19:86.

